# Cardiac electrophysiological remodeling associated with enhanced arrhythmia susceptibilty in a canine model of elite exercise

**DOI:** 10.1101/2022.07.13.499876

**Authors:** A. Polyák, L. Topal, J. Prorok, N. Tóth, Zs. Kohajda, Sz. Déri, V. Demeter-Haludka, P. Hegyi, V. Venglovecz, A. Sarusi, G. Ágoston, Z. Husti, N. Zombori-Tóth, P. Gazdag, J. Szlovák, T. Árpádffy-Lovas, M. Naveed, N. Jost, L. Virág, N. Nagy, I. Baczkó, A. S. Farkas, A. Varró

## Abstract

The health benefits of regular physical exercise are well known. Even so, there is increasing evidence that the exercise regimes of elite athletes can evoke cardiac arrhythmias including ventricular fibrillation and even sudden cardiac death (SCD). The mechanism of exercise-induced arrhythmia and SCD is poorly understood. While some studies after endurance training have been performed in small animals these have limited translation value.

Here, we show that chronic training in a canine model (12 sedentary and 12 trained dogs) that mimics the regime of elite athletes induces electrophysiological remodeling (measured by ECG, patch-clamp and immunocytochemical techniques) resulting in increases of both the trigger and the substrate for ventricular arrhythmias. Thus, 4 months sustained training lengthened ventricular repolarization (QT_c_: 213.6±2.8 ms vs. 237.1±3.4 ms, n=12; APD_90_: 370.1±32.7 ms vs. 472.8±29.6 ms, n=25 vs. 29), decreased transient outward potassium current (8.8±0.9 pA/pF vs. 6.4±0.5 pA/pF at 50 mV, n=42 vs. 54) and increased the short term variability of repolarization (17.5±4.0 ms vs. 29.5±3.8 ms, n=18 vs. 27). Left ventricular fibrosis and HCN4 protein expression were also enhanced. These changes were associated with enhanced ectopic activity (number of extrasystoles: 4/hour vs. 366/hour) *in vivo* and arrhythmia susceptibility (elicited ventricular fibrillation: 3 of 10 sedentary dogs vs. 6 of 10 trained dogs).

Our findings provide *in vivo*, cellular electrophysiological and molecular biological evidence for the enhanced susceptibility to ventricular arrhythmia in an experimental large animal model of endurance training.

## INTRODUCTION

The health benefits of regular physical exercise are well known [1, 2]. However, there is increasing evidence that chronic high level exercise in elite athletes can evoke cardiac arrhythmias including atrial fibrillation [3, 4] and even sudden cardiac death (SCD) [5]. Fortunately, sport-related SCD is rare although its incidence may often be under-estimated. Thus, SCD seems to be 2.8 times more frequent in elite athletes than in age-matched populations [6] who do not engage in sporting activity. It has to be mention that SCD incidence was found much higher than that in certain young athletes population such as male college basketball players [7]. In addition, in only few cases has cause of death been satisfactory established by autopsy findings; part of the remaining cases SCD has been attributed to ventricular fibrillation of ischemic origin. However, the latter explanation can be challenged because very often SCD in elite athletes does not occur during peak performance when oxygen demand is indeed very high in the myocardium. Instead, SCD occurs during warmup or after exercise or even at home during rest. Therefore, the cause and mechanism of SCD due to heavy chronic exercise should be also sought elsewhere.

It was recently reported that high levels of exercise in rats or mice induce electrophysiological remodeling resulting in atrial fibrillation [4, 8], sinus bradycardia [9] and atrioventricular node dysfunction [10]. Also, a recent study in rats after high intensity chronic exercise, found fibrosis and other cardiac dysfunction which were associated with enhanced vulnerability to arrhythmia at the supraventricular level [11]. The mechanism of SCD in human elite athletes is, for obvious reasons, very hard to study, while animal studies which were focused on AF after chronic endurance training were carried out in mice or in rats in which cardiac function eg. heart rate and repolarization properties, differs in important ways from human [4,8,11,12]. With the exception of one incomplete and preliminary study [13], and a very recent report on atrioventricular dysfunction in race horses [10], there have been no experimental studies of the subject in large animals which would have better translational value [14].

Accordingly, the aim of the present study was to determine the effect of 4 months sustained exercise on cardiac remodeling, and possible arrhythmia susceptibility, in a canine model that better reflects human physiology and pathophysiology.

## METHODS

### General methods

Animal maintenance and research were conducted in accordance with the National Institutes of Health Guide for the Care and Use of Laboratory Animals. All procedures using animals were approved by the Ethical Committee for the Protection of Animals in Research of the University of Szeged, Szeged, Hungary (approval numbers: I-74-15-2017 and I-74-24-2017) and by the Department of Animal Health and Food Control of the Ministry of Agriculture and Rural Development (authority approval numbers XIII/3330/2017 and XIII/3331/2017) and conformed to the rules and principles of the 2010/63/EU Directive.

### Experimental protocol

Before starting the experiments animals were conditioned to the exercise protocol. Those animals (2 dogs), which did not voluntarily cooperated properly were excluded from the further experiments. After the three weeks conditioning periods beagle dogs (purchased from a certified experimental animal breeder, Ásotthalom, Hungary; breeder authority approval number: XXXV/2018 by the Department of Animal Health and Food Control of the Ministry of Agriculture and Rural Development, Hungary), either sex, weighing 9–15 kg, were randomized into sedentary (SED, n=12) or trained (TRN, n=12) groups. All animals were 12 months old at the beginning of the training. Running sessions were performed on a special dog treadmill system (Dogrunner K9 Racer Treadmill, Dendermonde, Belgium) with controllable gradient and speed intensity. TRN animals underwent a 16-week-long training period, while SED group did not participate in the training. The protocol started with a 2-week warm-up period, thereafter animals were trained for 5 days a week with 2×90 minutes at speed 12-18 km·h −1 and with 2×50 minutes interval running at speeds of 4 and 22 km·h −1 a day for 16 weeks. A rest period was regularly applied to maintain appropriate hydration, but the total interruption did not exceed 1.5 hours in total. The training intensity was maintained with the use of 5% to 12% inclination. After each training session, dogs received portions of their preferred food as reward. The training protocol was tested in preliminary experiments and set to the maximum level which could be performed without distress. No a priori sample size estimation from a power calculation was done for this study. As a result, we collected as much data as was possible given limitations on funding for data collection from chronic experiments in large animals. The capacity of our laboratories and of the dog treadmills limited the group size, as well. The different experimental procedures were performed in a blinded fashion: the investigators were not aware of the groups when performing experiments/analyses. However, due to the nature of the experimental settings in case of the ECG measurements, where recordings were made before (‘self-control’) and following the 16-week-long training on the same animals, blinding could not be undertaken.

### Echocardiography, morphometry and histology

Echocardiography was performed at 0 and 16 weeks of the training protocol. M-mode parasternal long axis view was applied using 11.5 MHz transducer (GE 10S-RS, GE Healthcare, Chicago, IL, USA), connected to an echocardiographic imaging unit (Vivid S5, GE Healthcare, Chicago, IL, USA). All parameters were analysed by an investigator in a randomised and blinded manner. Left ventricular internal diameter during diastole (LVEDD), thickness of the left ventricular posterior wall (LVPW) and interventricular septum (IVS) were measured in M-mode images. These parameters were also normalized to body weight (BW) or body surface area (BSA). Animals were euthanized with pentobarbital sodium (150 mg/kg, i.v.) following sedation (xylazine 1 mg/kg, i.v.). After the corneal reflex of each dog had disappeared, the heart of the animals was excised. Total cardiac mass was measured after heart removal, then the atria were removed from the hearts and ventricles were separately weighed to yield the calculate heart-weight-to-body-weight and ventricular-weight-to-body weight ratios. Ventricular and septal wall thicknesses were also measured with a digital caliper. Samples were taken from the atria, septum and ventricular free-wall for histology. Paraffin sections were stained with Crossmon-trichrome to identify collagen deposition. Semiquantitative analysis was performed by an independent pathologist who scored the degree of interstitial fibrosis as: 0=negative; 1=mild; 2=moderate.

### Electrocardiography and open chest arrhythmia provocation

In conscious dogs, ECGs were measured using precordial leads at 0 and at 12 or 16 weeks. The methods have been described by Polyak *et al.* [13]. ECGs were recorded simultaneously with National Instruments data acquisition hardware (PC card, National Instruments, Austin, TX., U.S.A.) and SPEL Advanced Haemosys software (version 3.26, Experimetria Ltd. and Logirex Software Laboratory, Budapest, Hungary). RR, PQ, QRS, and QT intervals were measured by manual positioning on screen markers of 40 consecutive sinus beats at the 10th minute after initiation of the recording, then mean values were calculated. Heart rate was calculated from the RR interval. As QT interval is influenced by the heart rate, baseline data for ventricular heart rates and QT intervals were used to determine the relationship between the RR interval and the QT interval in sinus rhythm according to Kui *et al.* [15]. Simple linear regression revealed a positive correlation between QT and RR intervals (QT = 0.045RR + 186.9). The equations were rearranged to allow the calculation of the rate-corrected QT interval at an RR interval of 528 ms (i.e. a ventricular rate of 118 beats·min−1) using the formula QTc_x_ = QT_x_-0.045 (RR_(x−1)_-528). With these equations, plotting QTc against the corresponding RR interval produces a regression line with a slope of zero, indicating that these corrections remove the influence of heart rate.

Beat-to-beat variability and instability (BVI) parameters of the RR and QT intervals – e.g. the ‘root mean square of the successive differences’ (rmsSD) and the ‘short-term variability’ of the QT intervals (STV-QT) – were derived from 40 consecutive sinus beats as described previously [13].

Open chest arrhythmia provocation procedures were performed by four consecutive times of burst pacing (800/min equal to 13.3 Hz, 3× threshold in voltage, lasting for 1-3-6-9 seconds, were performed to induce ventricular fibrillation. The occurrence and manifestation time of VFs were compared.

### Conventional microelectrode techniques

Action potentials were recorded in left ventricular papillary muscle preparations obtained from the hearts of the exercised and sedentary dogs using the conventional microelectrode techniques previously described in detail [16]. Briefly, dogs were euthanized with pentobarbital sodium as described before. The animals also received an intravenous injection of 400 IU/kg heparin. Preparations were individually mounted in a tissue chamber with a volume of 50 ml. During experiments, modified Locke’s solution was used containing (mM): NaCl 128.3, KCl 4, CaCl_2_ 1.8, MgCl_2_ 0.42, NaHCO_3_ 21.4, and glucose 10. The pH of this solution was set between 7.35 and 7.4 when gassed with 95% O_2_ and 5% CO_2_ at 37°C. Each preparation was stimulated through a pair of platinum electrodes in contact with the preparation at a constant basic cycle length of 1000 ms. Transmembrane potentials were recorded after 60 min equilibrium time after mounting using conventional glass microelectrodes, filled with 3 M KCl The measurements where the resting membrane potential of the recorded action potential was more positive than −70 mV and/or the action potential amplitude was less than 90 mV were excluded from the analyses.

### Patch-clamp measurements

Dogs were euthanized with pentobarbital sodium as described before. The animals also received intravenous injection of 400 IU/kg heparin. Ventricular myocytes were enzymatically dissociated as previously described in detail [17]. One drop of cell suspension was placed in a transparent recording chamber mounted on the stage of an inverted microscope (Olympus IX51, Olympus, Tokyo, Japan), and individual myocytes were allowed to settle and adhere to the chamber bottom for at least 5-10 min before superfusion was initiated and maintained by gravity. Only rod-shaped cells with clear striations were used. HEPES-buffered Tyrode’s solution (composition in mM: NaCl 144, NaH_2_PO_4_ 0.4, KCl 4.0, CaCl_2_ 1.8, MgSO_4_ 0.53, glucose 5.5 and HEPES 5.0, at pH of 7.4) served as the normal superfusate.

Micropipettes were fabricated from borosilicate glass capillaries (Science Products GmbH, Hofheim, Germany), using a P-97 Flaming/Brown micropipette puller (Sutter Co, Novato, CA, USA), and had a resistance of 1.5-2.5 MOhm when filled with pipette solution. The membrane currents were recorded with Axopatch-200B amplifiers (Molecular Devices, Sunnyvale, CA, USA) by means of the whole-cell configuration of the patch-clamp technique. The membrane currents were digitized with 250 kHz analogue to digital converters (Digidata 1440A, Molecular Devices, Sunnyvale, CA, USA) under software control (pClamp 10, Molecular Devices, Sunnyvale, CA, USA). All patch-clamp experiments were carried out at 37 °C.

### Measurement of L-type calcium current

The L-type calcium current (I_CaL_) was recorded in HEPES-buffered Tyrode’s solution supplemented with 3 mM 4-aminopyridine. A special solution was used to fill the micropipettes (composition in mM: CsCl 125, TEACl 20, MgATP 5, EGTA 10, HEPES 10, pH was adjusted to 7.2 by CsOH).

### Measurement of potassium currents

The inward rectifier (I_K1_), transient outward (I_to_), rapid (I_Kr_) and slow (I_Ks_) delayed rectifier potassium currents were recorded in HEPES-buffered Tyrode’s solution. The composition of the pipette solution (mM) was: KOH 110, KCl 40, K_2_ATP 5, MgCl_2_ 5, EGTA 5, HEPES 10 (pH was adjusted to 7.2 by aspartic acid). 1 µM nisoldipine was added to the bath solution to block I_CaL_. When I_Kr_ was recorded I_Ks_ was inhibited by using the selective I_Ks_ blocker HMR 1556 (0.5 µM). During I_Ks_ measurements, I_Kr_ was blocked by 0.1 µM dofetilide and the bath solution contained 0.1 µM forskolin.

### Measurement of late sodium current

The sodium current is activated by 2 s long depolarizing voltage pulses to −20 mV from the holding potential of −120 mV with pulsing cycle length of 5 s. After 5 - 7 min incubation with the drug, the external solution was replaced by that containing 20 µM TTX. TTX at this concentration completely blocks the late sodium current (I_NaL_). The external solution was HEPES-buffered Tyrode’s solution supplemented with 1 µM nisoldipine, 0.5 µM HMR-1556 and 0.1 µM dofetilide in order to block I_CaL_, I_Ks_ and I_Kr_ currents. The composition of the pipette solution (in mM) was: CsCl 125, TEACl 20, MgATP 5, EGTA 10, HEPES 10, pH was adjusted to 7.2 by CsOH).

### Measurement of NCX current

For the measurement of the Na^+^/Ca^2+^ exchanger current (I_NCX_), the method of Hobai *et al.* [18] was applied. Accordingly, the NCX current is defined as Ni^2+^-sensitive current and measured in a special K^+^-free solution (composition in mM: NaCl 135, CsCl 10, CaCl_2_ 1, MgCl_2_ 1, BaCl_2_ 0.2, NaH_2_PO_4_ 0.33, TEACl 10, HEPES 10, glucose 10 and ouabain 20 µM, nisoldipine 1 µM, and lidocaine 50 µM, at pH 7.4) as described earlier in detail [17]. The pipette solution used for recording I_NCX_ contained (in mM) CsOH 140, aspartic acid 75, TEACl 20, MgATP 5, HEPES 10, NaCl 20, EGTA 20 and CaCl_2_ 10, pH adjusted to 7.2 with CsOH.

### Measurements of single cell action potentials

The perforated patch-clamp technique was used to measure action potentials from isolated left ventricular myocytes from both trained and sedentary animals. The membrane potential was recorded in current clamp configuration. The myocytes were paced with a rapid rectangular pulse (from 0 to 180 mV, 5 ms) at a frequency of 1 Hz to elicit the action potential. A normal Tyrode solution was used as the extracellular solution containing (in mM): 144 NaCl, 0.4 NaH_2_PO_4_, 4 KCl, 0.53 MgSO_4_, 1.8 CaCl_2_, 5.5 glucose and 5 HEPES, titrated to pH 7.4. The patch pipette solution contained (in mM): 120 K-gluconate, 2.5 NaCl, 2.5 MgATP, 2.5 Na_2_ATP, 5 HEPES, 20 KCl, titrated to pH 7.2 with KOH. 50 µM β-escin was added to the pipette solution to achieve the membrane patch perforation. Membrane voltage was obtained by using an Axoclamp 1-D amplifier (Molecular Devices, Sunnyvale, CA, USA) connected to a Digidata 1440A (Molecular Devices, Sunnyvale, CA, USA) analogue-digital converter. The membrane voltage was recorded by Clampex 10.0 (Molecular Devices, Sunnyvale, CA, USA). At least 60 beats were recorded, and the action potential duration was measured at 90% repolarization (APD_90_). The short term APD variability was calculated by analysing 30 consecutive action potentials.

### Western blot analysis of KChIP2 and Kv4.3

Membrane fractions were isolated from myocardial samples (n=24) taken from left ventricular wall using the method described previously [19]. Protein concentrations were determined by the Lowry method and 20 µg of each sample was then separated on 8% polyacrylamide gels and transferred to PVDF membrane. The membrane was blocked with 2.5% non-fat milk for 1 hour at room temperature and immunolabelled overnight at 4°C with anti-KChIP2 (Alomone, #APC-142, RRID:AB_2756744) and anti-Kv4.3 (Alomone, #APC-017, RRID:AB_2040178) primary antibody diluted to 1:1000. This was followed by 1 hour incubation with Goat anti-Rabbit IgG-HRP (SouthernBiotech, 4030-05, RRID:AB_2687483) secondary antibody in a dilution of 1:8000. Band densities were detected with ECL Prime Western Blotting Detection Reagent (GE Healthcare) and a ChemiDoc Imaging System (Bio-Rad). Equal loading was verified by GAPDH labelling (ThermoFisher, PA1-988, RRID:AB_2107310). Pixel intensities of each band were measured using ImageJ software. Three parallel Western blots were performed for the statistical analysis.

### Immunocytochemistry of KChIP2, Kv4.3, HCN1, HCN2 and HCN4

Cardiomyocytes were isolated from left ventricular tissue (*n*= 6 trained and 6 sedentary dogs), then fixed on glass coverslips with acetone [20]. Before immunolabelling, samples were rehydrated with calcium-free phosphate-buffered saline (PBS) and blocked for 1 hour with PBST (PBS with 0.01% Tween) containing 2.5% BSA (bovine serum albumin) at room temperature. After the incubation period, cells were labelled overnight at 4°C with anti-KChIP2 (Alomone, #APC-142, RRID:AB_2756744), anti-Kv4.3 (Alomone, #APC-017, RRID:AB_2040178), anti-HCN1 (Alomone, #APC-056, RRID:AB_2039900), anti-HCN2 (Alomone, #APC-030, RRID:AB_2313726) and anti-HCN4 (Alomone, #APC-052, RRID:AB_2039906) primary antibody diluted to 1:50. The next day, cells were incubated with goat anti-rabbit IgG Alexa Fluor 488 (ThermoFisher, A-11034, RRID:AB_2576217) secondary antibody (dilution: 1:500, ThermoFisher). Fluorescent images were captured by an LSM 880 (Zeiss) laser scanning confocal microscope. Images were quantitatively analyzed by the ImageJ software. Control samples were incubated only with secondary antibody.

### Statistics

IBM SPSS Statistics V25 and Microsoft Excel (Microsoft Office Professional Plus 2016) software packages were used for statistical analysis. Continuous data were expressed as mean ± standard error of the mean (S.E.M.). Each figure stipulates the number of observations made (“n”), represents the biological replication of the experiment. The “n” refer to the number of dogs except in action potential and patch clamp experiments and in immunocytochemistry measurements where it refer to the number of preparations/cells followed by the number of dogs where preparations/cells were obtained from. Paired and unpaired Student’s t-test were applied to estimate whether there was a statistically significant difference between the means in self-control or independent group arrangements, respectively. Data were considered statistically significant when p < 0.05.

## RESULTS

### Sustained exercise induced cardiac hypertrophy and fibrosis

The 16 weeks endurance training in dogs resulted significantly greater end diastolic diameter (LVEDD) in the left ventricle compared to sedentary controls or to exercised animals before the training protocol (TABLE 1) measured by *in vivo* by echocardiography. The calculated left ventricular mass (LVM), the thickness of the left ventricular posterior wall (LVPW) and the interventricular septum (IV) were also increased in exercised dogs compared to sedentary controls or pre-training controls.

**Table 1.** The effect of exercise training on echocardiographic cardiac dimensions, autopsy findings, and the ECG parameters in dog heart. Echocardiographic values were measured before (at 0th week, control measurements) and after (at 16th week) the training protocol. IVS, Interventricular septum; BW, body weight; LVPW, Left ventricular posterior wall thickness; LVEDD, Left ventricular end diastolic dimension; BSA, body surface area; LVM, left ventricular mass; LVMi, Left ventricular mass index, calculated by dividing the LVM by the BSA; EDV, end diastolic volumen; SV, stroke volumen, Cardiac index, calculated by dividing the cardiac output by the BSA; LAV, left atrial volumen; LAVi, left atrial volumen index, calculated by dividing the LAV by the BSA. *P<0.05 vs. Trained 0 week. All values are means±SEM. **Table 1–Source Data 1** The effect of exercise training on echocardiographic cardiac dimensions. **Table 1–Source Data 2** The effect of exercise training on ECG parameters.

| Echocardiographic dimensions | DOG |  |  |  |
| --- | --- | --- | --- | --- |
|  | Before the training protocol (0<sup>th</sup> week) |  | After the training protocol (16<sup>th</sup> week) |  |
|  | Sedentary (n=12) | Trained (n=12) | Sedentary (n=12) | Trained (n=12) |
| IVS, mm | 7.1±0.3 | 6.8±0.2 | 7.4±0.3 | 8.1±0.2 |
| IVS/BW, mm | 0.6±0.03 | 0.5±0.03 | 0.6±0.03 | 0.7±0.03 |
| LVPW, mm | 7.1±0.2 | 7.0±0.2 | 7.4±0.3 | 7.6±0.3 |
| LVPW/BW, mm | 0.6±0.03 | 0.6±0.03 | 0.6±0.03 | 0.7±0.04 |
| LVEDD, mm | 28.7±0.7 | 29.0±1.0 | 30.4±0.7 | 32.0±0.7 |
| LVEDD/BSA, mm | 51.7±1.4 | 52.7±1.0 | 55.3±2.0 | 63.5±1.3 |
| LVM, g | 46.7±3.4 | 45.1±2.6 | 54.1±3.9 | 63.6±2.8 |
| LVMi, g | 83.6±5.6 | 81.2±2.8 | 97.7±6.4 | 125.8±4.3 |
| EDV, ml | 32.3±2.0 | 32.6±2.5 | 37.9±2.2 | 40.6±1.7 |
| EDV/BSA, ml | 57.5±2.5 | 58.4±2.9 | 68.4±3.4 | 80.3±2.3 |
| SV, ml | 26.2±1.9 | 23.1±2.0 | 26.5±1.6 | 30.4±1.4 |
| Cardiac index | 5.1±0.5 | 4.2±0.3 | 4.3±0.3 | 5.5±0.4 |
| LAV, ml | 9.0±0.9 | 8.9±0.6 | 10.4±0.9 | 11.4±1.4 |
| LAVi, ml | 16.1±1.4 | 16.0±0.8 | 18.7±1.3 | 22.4±2.3 |
| Autopsy findings |  |  |  |  |
| LVM, g |  |  | 79.3±4.4 | 83.3±4.8 |
| LVMi, g |  |  | 148.5±5.4 | 167.2±5.7 |
| IVS, mm |  |  | 3.6±0.6 | 4.3±0.3 |
| IVS/BW, mm |  |  | 0.3±0.04 | 0.4±0.03 |
| LVPW, mm |  |  | 2.5±0.4 | 3.4±0.3 |
| LVPW/BW, mm |  |  | 0.2±0.03 | 0.3±0.02 |
| ECG parameters |  |  |  |  |
| RR (ms) | 516.1±31.5 | 544.3±22.8 | 587.5±28.5 | 828.9±63.8* |
| PQ (ms) | 103.2± 2.0 | 98.3± 2.9 | 102.8± 3.2 | 110.7± 3.6* |
| QRS (ms) | 59.6± 1.7 | 60.5± 2.4 | 56.3± 2.6 | 70.8± 1.6* |
| QT (ms) | 218.7± 5.7 | 215.9± 2.9 | 223.0± 6.4 | 251.3± 3.2* |
| QTc (ms) | 216.0± 4.7 | 213.6± 2.8 | 217.7± 4.5 | 237.1± 3.4* |
| STV-QT (ms) | 2.6± 0.2 | 2.5± 0.2 | 2.6± 0.2 | 3.6± 0.4* |

Autopsy findings in hearts obtained from exercised and sedentary dogs showed cardiac hypertrophy reflected an increased left ventricular mass (LVM) and thickening of the intraventricular septum (IVS) and left ventricular posterior wall (LVPW) with exercise (TABLE 1). In addition, some degree of enhanced fibrosis was also present in the hearts of exercised dogs compared with sedentary controls. (FIGURE 1)

**FIGURE 1.**
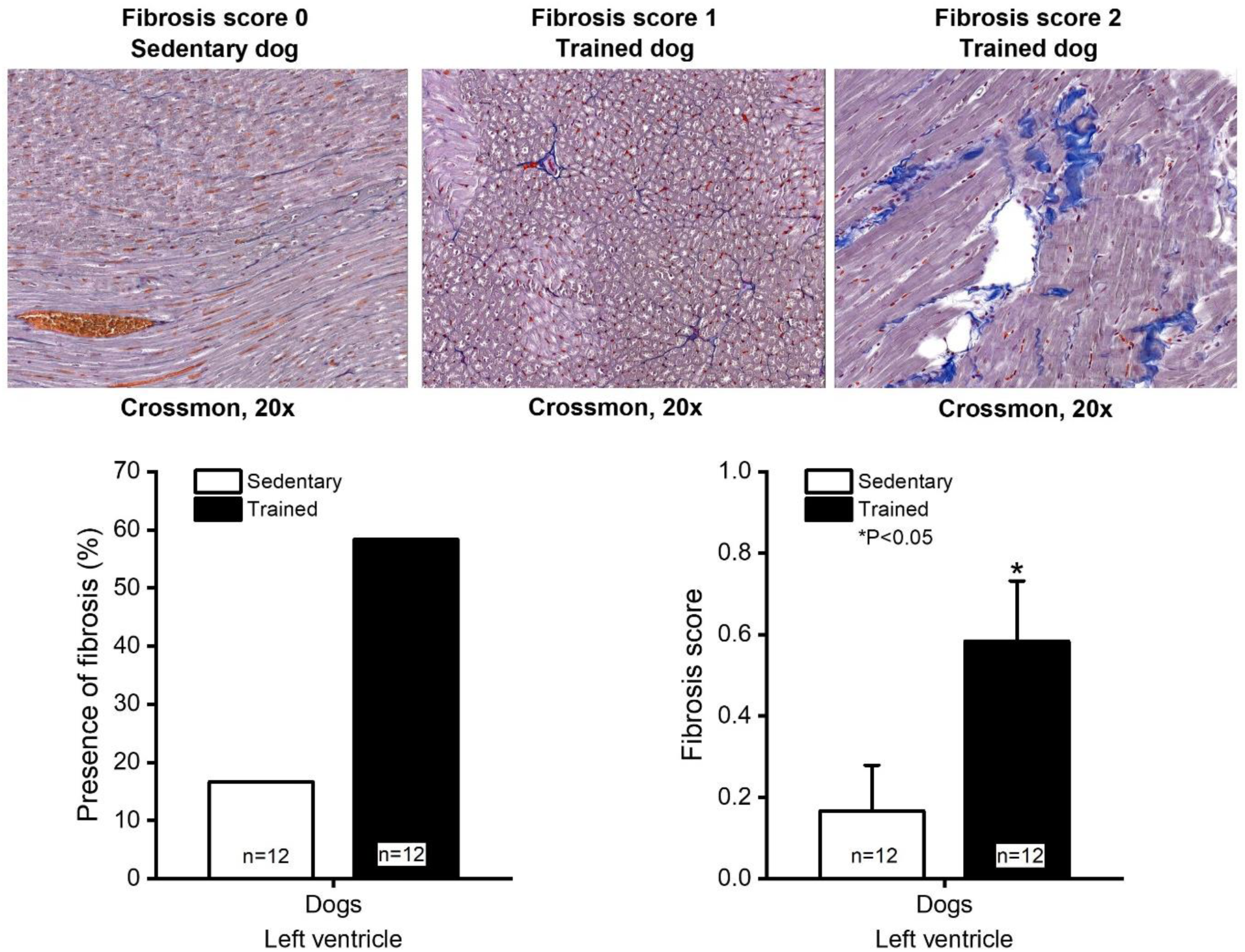
Enhanced fibrosis in the hearts of exercised dogs. Upper panels, representative histology of connective tissue visualized using Crossmon’s trichrome staining from sedentary control (fibrosis score 0), trained dogs (fibrosis score 1 and fibrosis score 2). Lower panels, bar charts of fibrosis (*left*) and the fibrosis score (*right*) in sedentary (n=12 dogs) and trained (n=12) dogs. Semiquantitative analysis was performed to score the degree of the interstitial fibrosis with the following criteria: 0=negative; 1=mild; 2=moderate. **Figure 1–Source Data 1** The effect of exercise training on presence of fibrosis. **Figure 1–Source Data 2** The effect of exercise training on fibrosis score.

### Effect of sustained training on the heart rate

Sustained training caused significant bradycardia and increase of the heart rate variability (rmsSD) in exercised dogs compared with sedentary controls (FIGURE 2 A). To study heart rate changes independent of possible alterations in vagal tone, heart rate was also studied in spontaneously beating isolated right atrial tissue preparations obtained from control or exercised dogs. The latter showed slower spontaneous frequency than sedentary controls (FIGURE 2 B) further suggesting that the bradycardia observed in exercised dogs is not entirely due to enhancement of the vagal tone.

**Figure 2.**
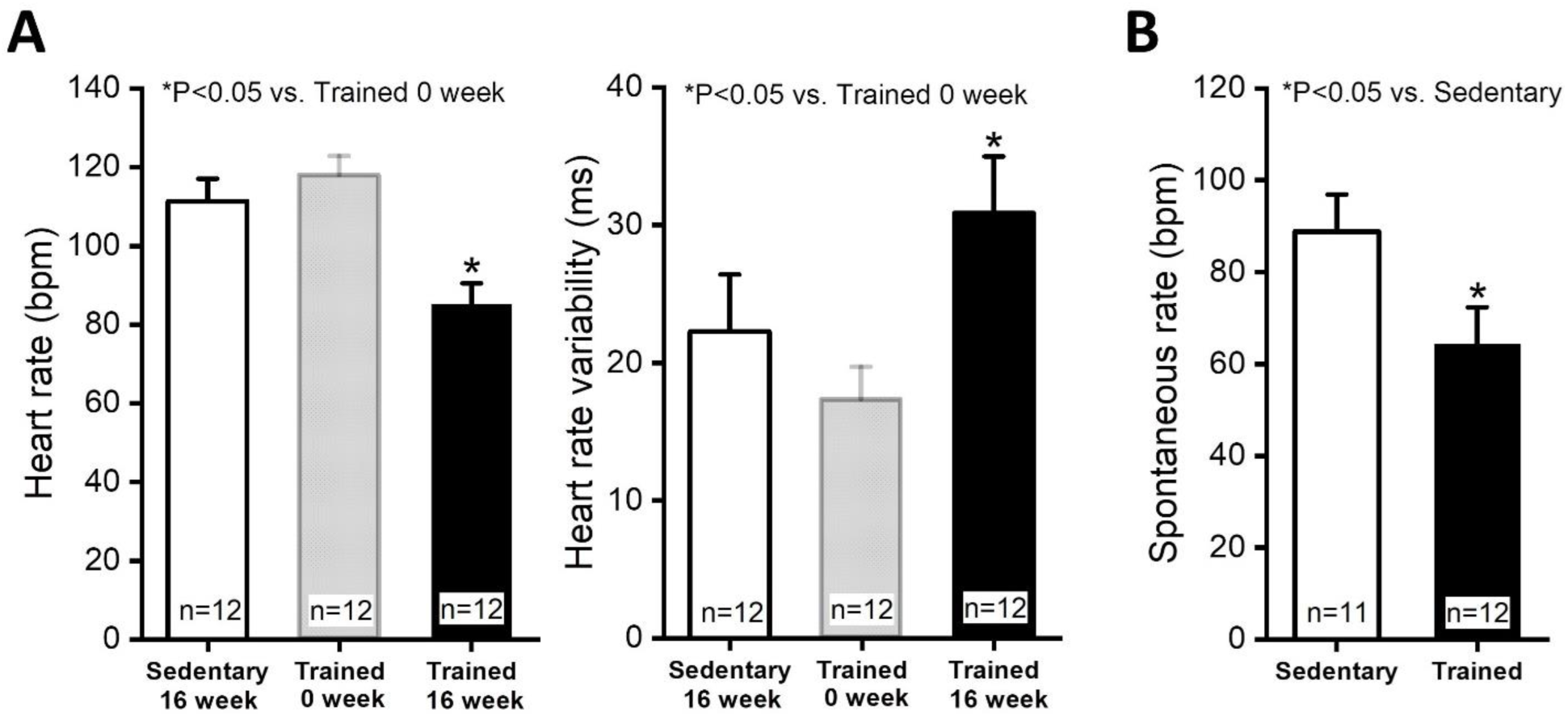
Effect of sustained training on heart rate in conscious dogs and on the spontaneous frequency of isolated right atrial tissue preparations. Panel A, heart rate (*left*) and the heart rate variability – rmsSD – (*right*) of conscious dogs in the sedentary (16 weeks – n=12 dogs), pretrained (0 week – before the chronic endurance training initiated) and trained groups (16 weeks – after chronic endurance training – n=12 dogs). Panel B, spontaneous frequency of isolated right atrial tissue preparations obtained from sedentary (n=11 dogs) and trained (n=12) dogs is indicated by the bar diagram. **Figure 2–Source Data 1** Effect of sustained training on heart rate in conscious dogs. **Figure 2–Source Data 2** Effect of sustained training on heart rate variability in conscious dogs. **Figure 2–Source Data 3** Effect of sustained training on the spontaneous frequency of isolated right atrial tissue preparations.

### ECG changes and enhanced proarrhythmic response due to sustained training

The 16 week sustained training lengthened RR, PQ, QT, QTc intervals and widened QRS significantly in conscious dogs (TABLE 1). The lengthened QT interval was also associated with significantly enhanced STV-QT interval reflecting elevated dispersion of repolarization measured after the completion of training protocol in exercised animals compared to sedentary controls (TABLE 1).

In 12 conscious sedentary dogs only 4 ventricular ectopic beats were observed in 2 animals during a 3 x 20 min resting periods. In the exercised 12 dogs however in 50 % of the animals, 366 ventricular ectopic beats were noticed (FIGURE 3 A).

**Figure 3.**
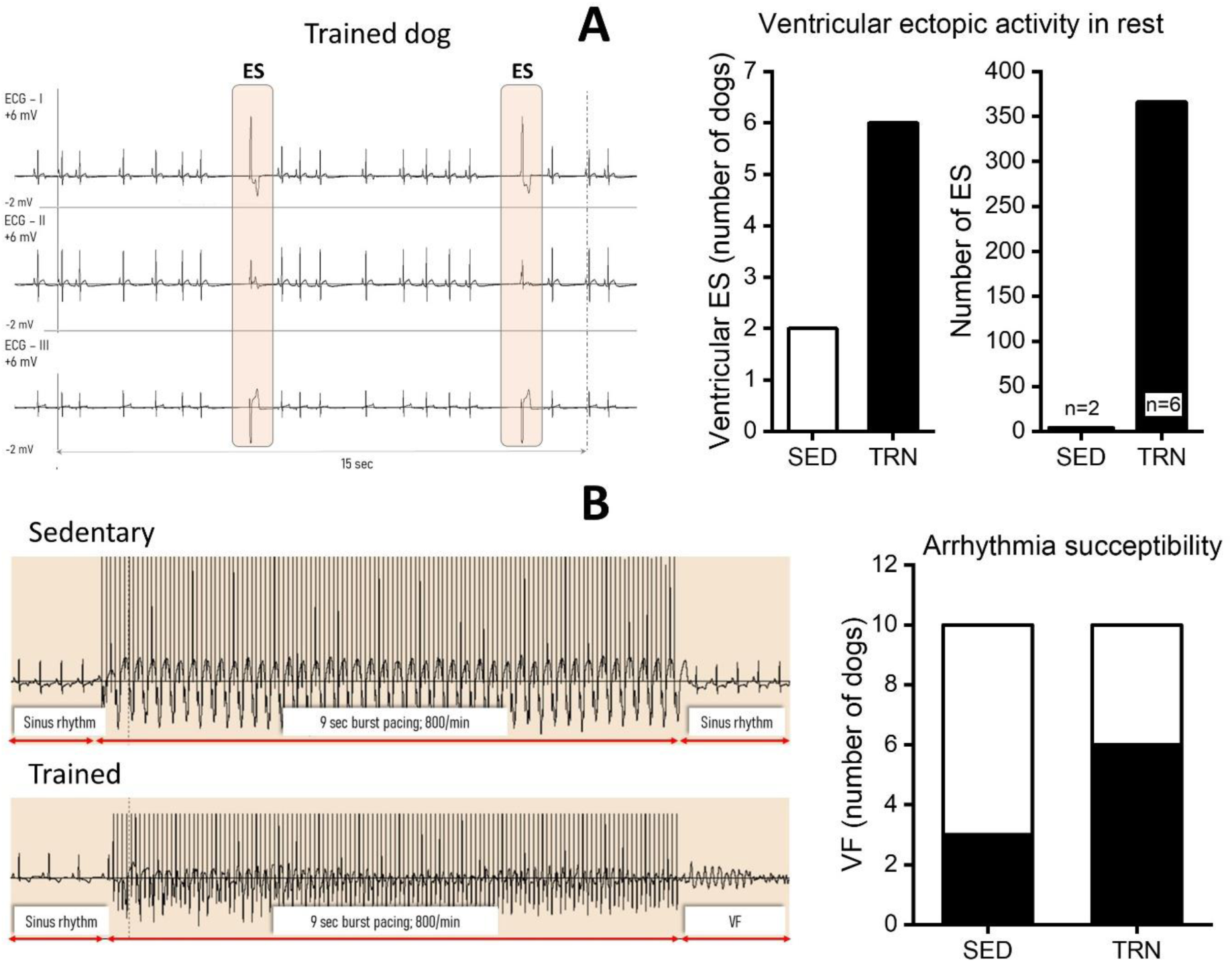
Incidence of ventricular extrasystoles and ventricular fibrillation in trained and sedentary dogs. Panel **A**: At rest, ventricular extrasystoles (ES) were observed in 6 of the trained animals (TRN) over a 3 x 20 min periods, whereas ES occurred in only two of the sedentary (SED) dogs. The number of extrasystoles was also significantly higher in trained animals (n=6 dogs) compared to untrained dogs (n=2). Panel **B**: Ventricular fibrillation (VF) occurred in 6 trained dogs as a result of ventricular burst arrhythmia provocation, whereas only 3 of the sedentary animals showed VF. Representative ECG recordings are shown on the left side of the figure. **Figure 3–Source Data 1** Incidence of ventricular extrasystoles in trained and sedentary dogs. **Figure 3–Source Data 2** The number of extrasystoles in trained and sedentary dogs. **Figure 3–Source Data 3** Incidence of ventricular fibrillation in trained and sedentary dogs.

In order to study the possible changes in arrhythmia susceptibility in open chest anaesthetized dogs, 800/s frequency electrical bursts were applied for 1-9 s. In these experiments TdP arrhythmias and consequent ventricular fibrillation (VF) was elicited in 6 of 10 exercised dogs compared to only 3 of 10 of the sedentary control dogs (FIGURE 3 B).

### Influence of sustained training on cardiac ventricular action potentials

Cardiac ventricular action potentials were measured by conventional microelectrode techniques in isolated left ventricular papillary muscle preparations representing subendocardial origin and also in single enzymatically isolated left ventricular myocytes representing midmyocardial origin obtained from control sedentary and exercised dogs (FIGURE 4). As FIGURE 4 shows, the cardiac action potential duration measured as 90 % repolarization (APD_90_) was not significantly different between the left ventricular papillary muscle preparations originating from exercised and sedentary dogs (FIGURE 4) but repolarization reflected as APD_90_ was significantly lengthened and APD-STV increased in the left ventricular myocytes isolated from exercised dogs compared to sedentary animals (FIGURE 4).

**Figure 4.**
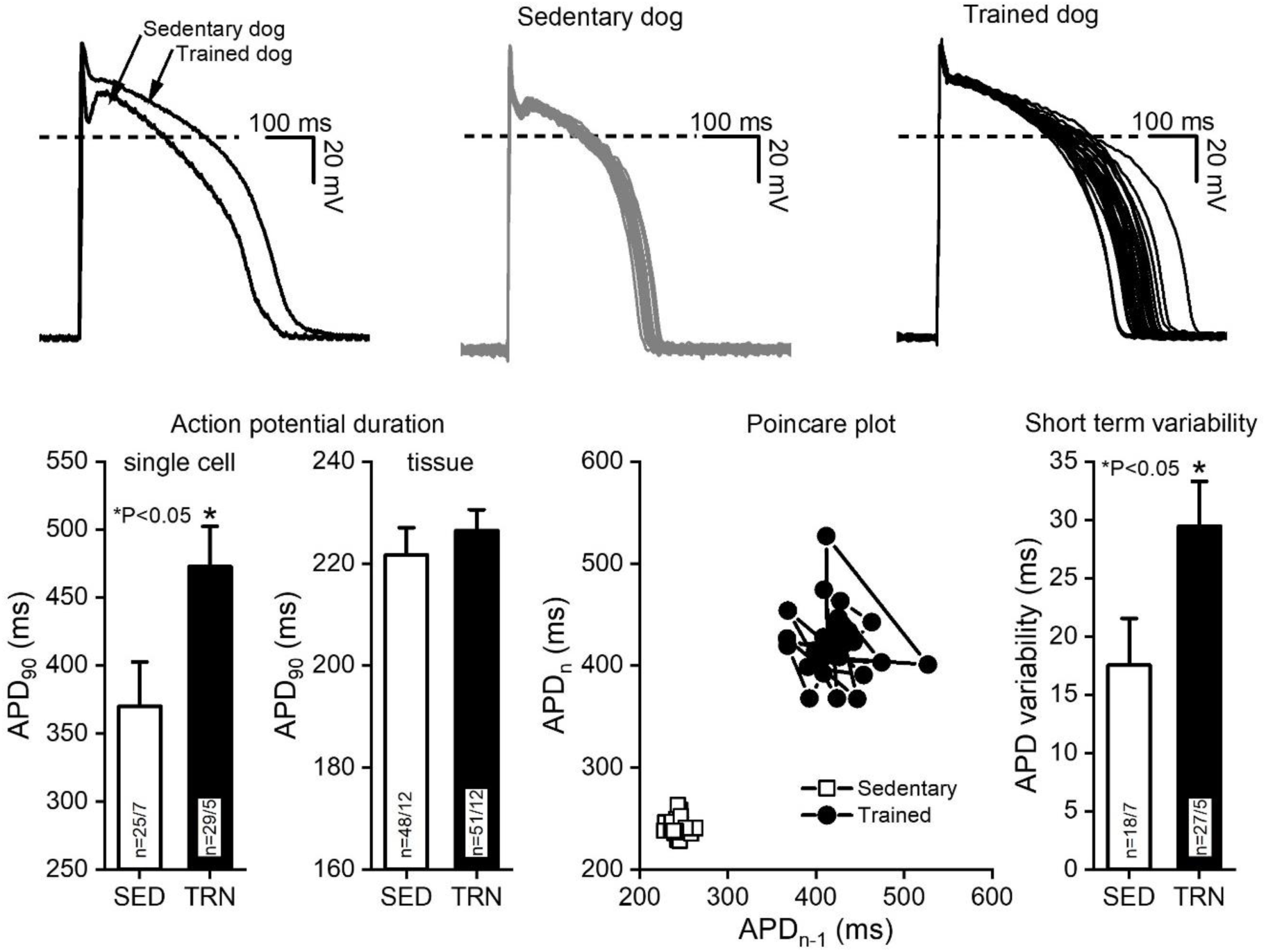
Effect of sustained training on the cardiac action potential duration and the short term variability in left ventricular preparations of control sedentary and exercised dogs. Upper panels, two representative action potential curves recorded from isolated left ventricular myocytes of a sedentary and a trained dog, respectively (*left*). *Centre* and *right,* 30 – 30 representative action potential curves indicating the variability of the action potential duration recorded from myocytes obtained from a sedentary and a trained dog, respectively. Lower panels, bar graphs showing action potential duration measured as 90 % repolarization (APD_90_) in sedentary (SED) and trained (TRN) dogs recorded from single myocytes (n=25 cells/7 dogs for sedentary and n=29 cells/5 dogs for trained groups) and from multicellular tissue preparations (n=48 preparations/12 dogs for sedentary and n=51 preparations/12 dogs for trained groups). Panels on the right side indicate the effect of chronic training on the short-term variability of the action potential duration in two representative Poincare plots (sedentary and trained) and in a bar graph (*right*). Action potentials were recorded from single left ventricular myocytes for these measurements obtained from sedentary (n=18 cells/7 dogs) and trained (n=27 cells/5) dogs. **Figure 4–Source Data 1** Effect of sustained training on the cardiac action potential duration (APD_90_) recorded from single myocytes. **Figure 4–Source Data 2** Effect of sustained training on the cardiac action potential duration (APD_90_) recorded from multicellular tissue preparations. **Figure 4–Source Data 3** Effect of sustained training on the short-term variability of the action potential duration.

### Possible influence of sustained training on the cardiac ventricular transmembrane currents

FIGURE 5 compares the transmembrane currents measured in enzymatically isolated left ventricular myocytes by the whole cell configuration of the patch clamp technique originating from the sedentary control and exercised dogs. As FIGURE 5 A shows I_to_ was significantly smaller in myocytes obtained from chronically trained dogs compared sedentary controls. There were, however, no significant differences in the current magnitude of the I_K1_, I_Kr_, I_Ks_, L-type I_Ca_, I_NaL_ and NCX currents (FIGURE 5 B,C,D,E,F,G).

**Figure 5.**
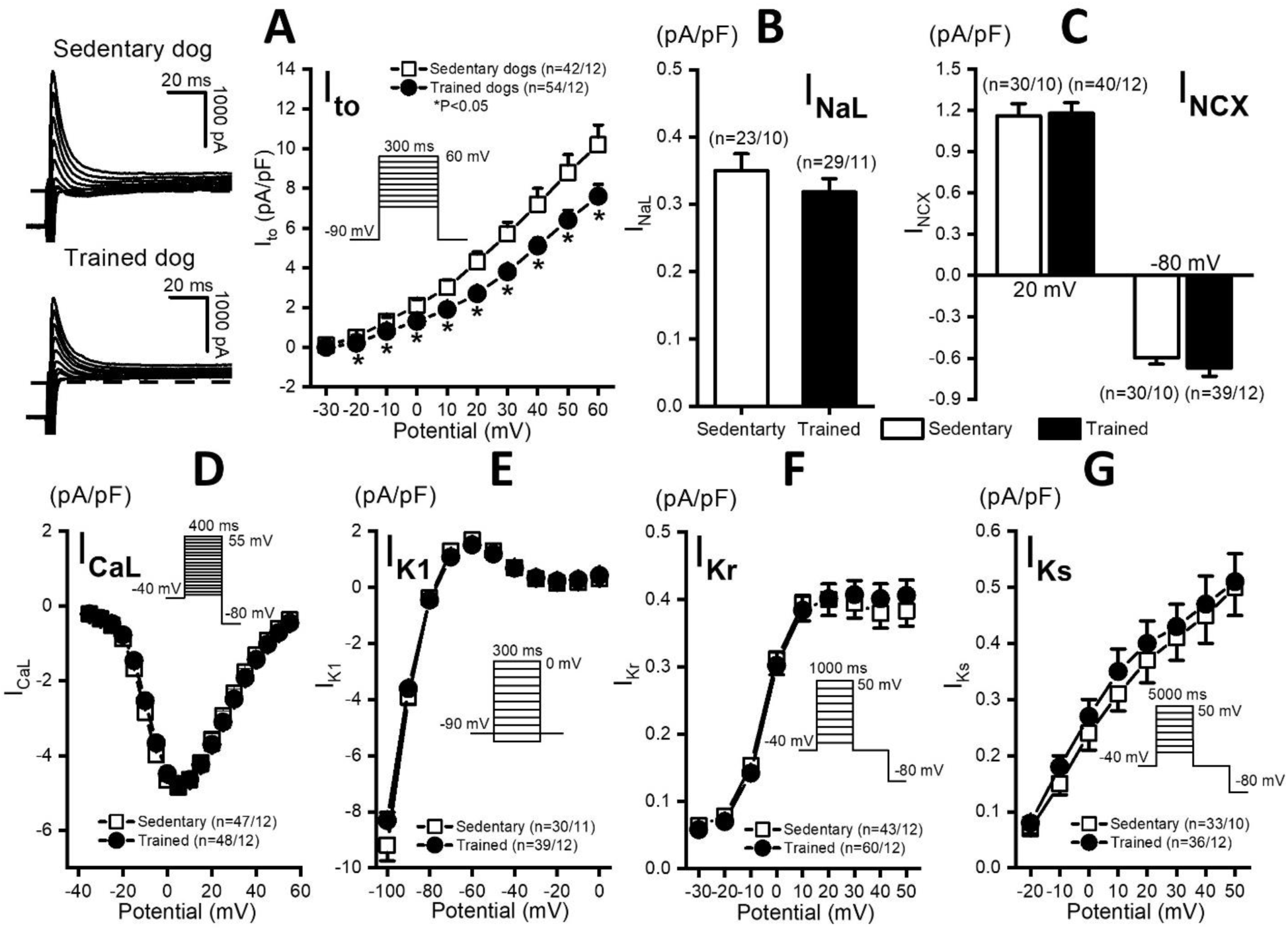
Effect of sustained training on various transmembrane ion currents in dog ventricular myocytes. Panel **A,** the effect of sustained training on transient outward potassium current (I_to_) showing original current records (*left*) and current-voltage relationships (*right*) for sedentary and trained individuals. Panels **B** and **C**, bars graphs indicate sustained training has no effect on the late Na^+^ current (I_NaL_) and on the Na^+^-Ca^2+^ exchange current (I_NCX_). Panels **D**, **E**, **F** and **G**, respectively, current-voltage relationships of L-type Ca^2+^ current (I_CaL_), inward rectifier K^+^ current (I_K1_), rapid (I_Kr_) and slow (I_Ks_) delayed rectifier K^+^ currents are similar for sedentary and trained individuals Insets show the voltage protocols. The “n” numbers refer to the number of isolated cells followed by the number of dogs where cells were obtained from. **Figure 5–Source Data 1** The effect of sustained training on the current-voltage relationship of transient outward potassium current. **Figure 5–Source Data 2** The effect of sustained training on late Na^+^ current. **Figure 5–Source Data 3** The effect of sustained training on Na^+^-Ca^2+^ exchange current. **Figure 5–Source Data 4** The effect of sustained training on the current-voltage relationship of L-type Ca^2+^ current. **Figure 5–Source Data 5** The effect of sustained training on the current-voltage relationship of inward rectifier K^+^ current. **Figure 5–Source Data 6** The effect of sustained training on the current-voltage relationship of rapid delayed rectifier K^+^ current. **Figure 5–Source Data 7** The effect of sustained training on the current-voltage relationship of slow delayed rectifier K^+^ current.

### Transmembrane Kv4.3 and KChiP2 protein abundance in exercised and sedentary dog hearts

The I_K1_, I_Kr_, I_Ks_, I_Ca_, I_NaL_ and NCX transmembrane current measurements showed no significant differences between sedentary and exercised dog hearts. The only transmembrane current which showed reduced amplitude was I_to_. In agreement with the transmembrane current data, significant APD lengthening was observed only in myocytes harvested from the left ventricular midmyocardial region where strong I_to_ can be expected [21, 22] but not in preparations of subendocardial left ventricular papillary muscle where relatively weak I_to_ was reported [22, 23]. Therefore, further studies of the molecular nature of the reduced I_to_ in exercised dog heart were performed. Specifically, the expression of Kv4.3 alfa and KChiP2 beta channel subunits, which are considered to be the most important channel proteins underlying I_to_, were studied by Western blot and immunocytochemical techniques. As FIGURE 6 indicates no significant difference of Kv4.3 and KChiP2 protein expression was found between the sedentary and exercised dog hearts suggesting that decrease of I_to_ is due to changes in other less well characterized accessory proteins or perhaps post translation changes in ion channel proteins.

**Figure 6.**
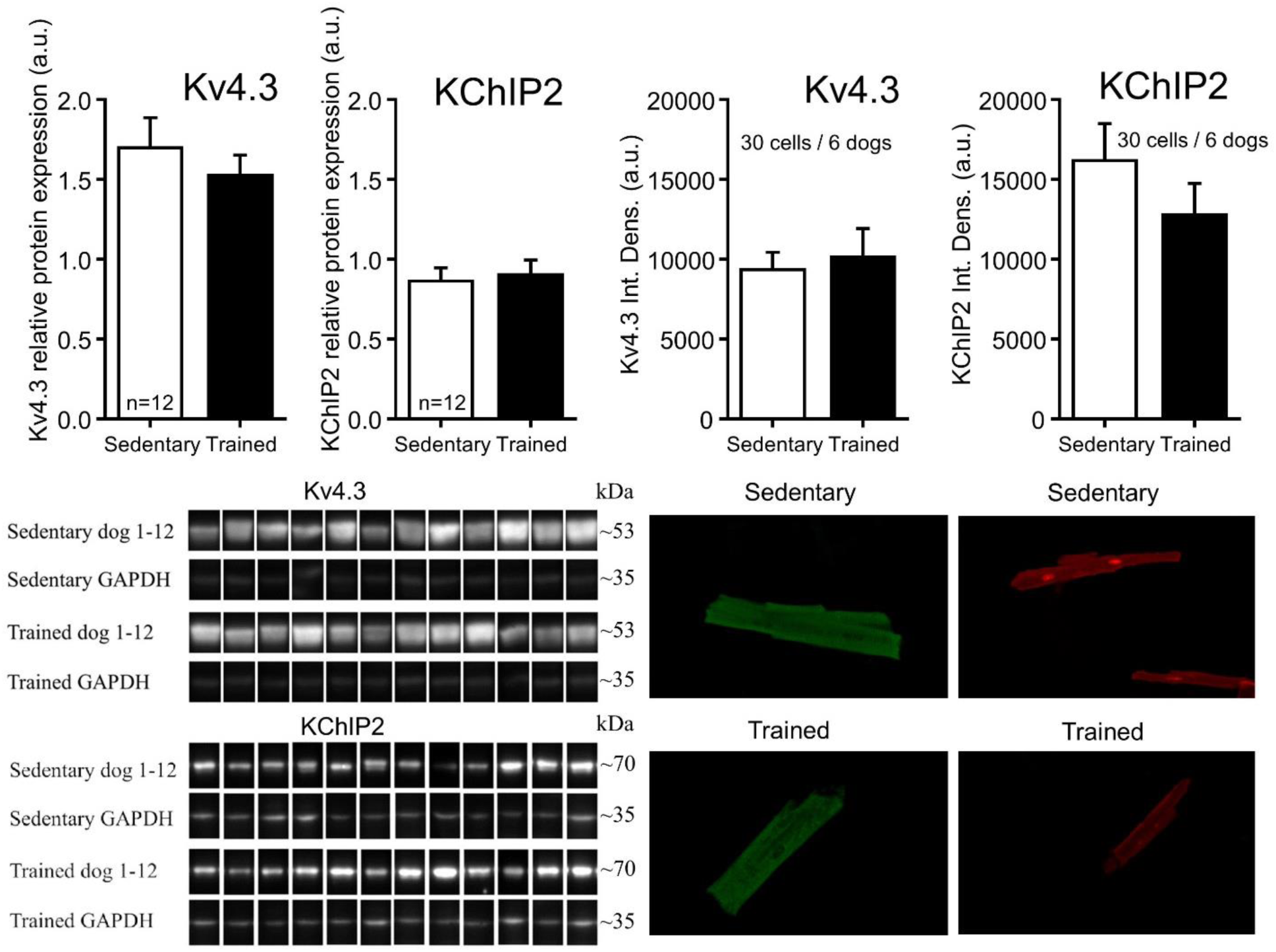
Protein expression and relative density of Kv4.3 and KChiP2 subunits determined by Western blot and immunocytochemical techniques in sedentary and trained dog. Two bar diagrams on the *left* show the relative protein expression of Kv4.3 and KChiP2 subunits determined by Western blot in sedentary (n=12 dogs) and trained dog (n=12) left ventricular samples. *Bottom* panels indicates the representative Kv4.3 and KChIP2 bands and their corresponding loading control (GAPDH). Two bar diagrams on the *right* panels show the relative density of dog cardiomyocytes with Kv4.3 and KChiP2 immunolabelling obtained from the sedentary (n=30 cells/6 dogs) and trained (n=30 cells/6 dogs) groups. Bottom panels on the *right* show original immunofluorescent images of dog cardiomyocytes with Kv4.3 and KChiP2 immunolabelling. **Figure 6–Source Data 1** Protein expression of Kv4.3 subunit determined by Western blot technique in sedentary and trained dogs. **Figure 6–Source Data 2** Protein expression of KChiP2 subunit determined by Western blot technique in sedentary and trained dogs. **Figure 6–Source Data 3** Relative density of Kv4.3 subunit determined by immunocytochemical technique in sedentary and trained dogs. **Figure 6–Source Data 4** Relative density of KChiP2 subunit determined by immunocytochemical technique in sedentary and trained dogs. **Figure 6–Source Data 5** Original, unedited membranes of western blots with the relevant bands clearly labelled. **Figure 6–Source Data 6** Original files of the full raw unedited membranes of western blots.

### Upregulation of HCN4 channels in the dog ventricle after sustained training

In myocytes obtained from exercised dog heart, immunohistochemistry showed enhanced HCN4 protein expression compared to sedentary dogs. There were no differences in expression of HCN1 and HCN2 proteins between the two groups (FIGURE 7).

**Figure 7.**
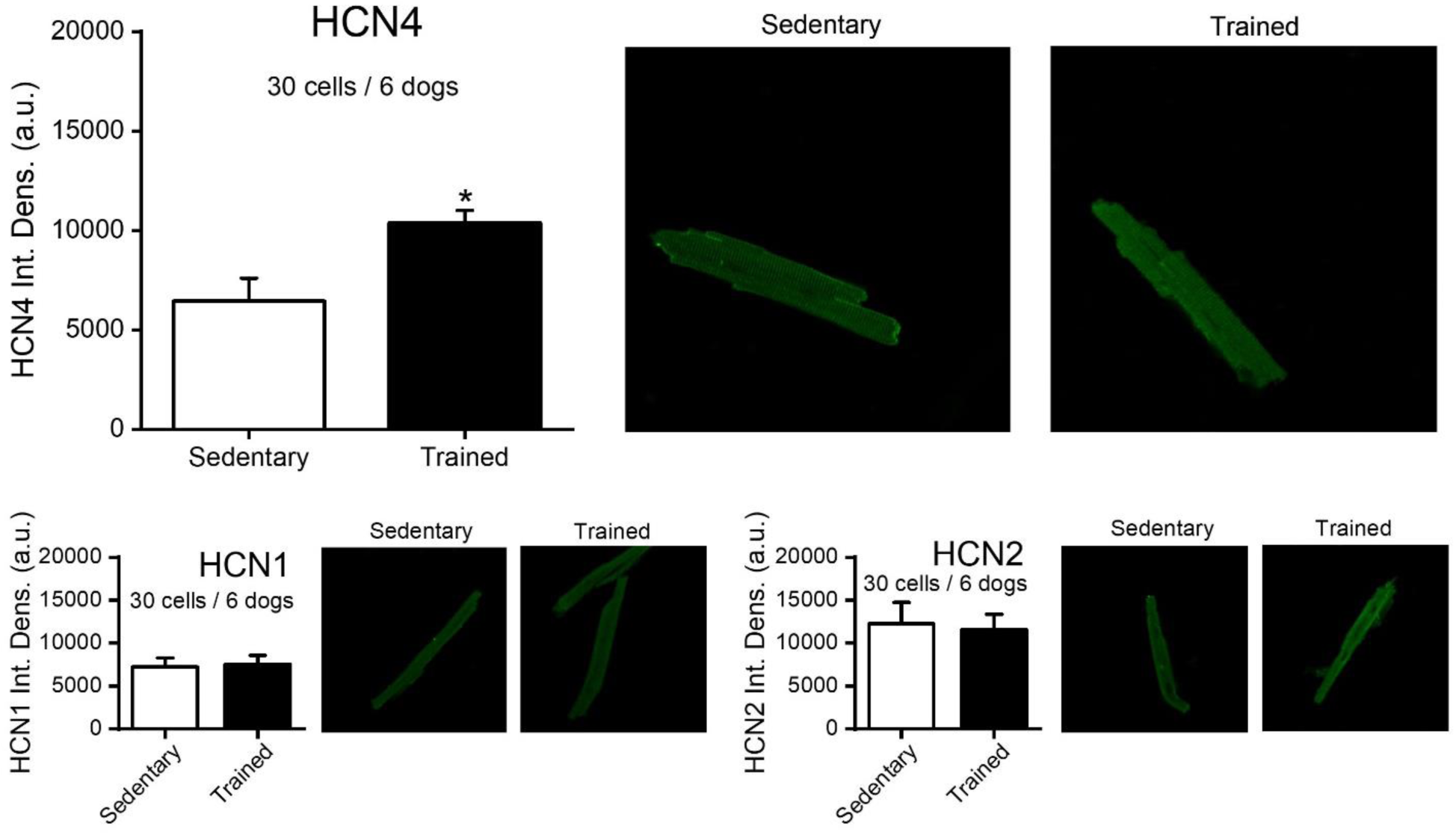
Effect of chronic training on HCN1, KCN2 and HCN4 protein expression determined by immunocytochemical techniques in sedentary and trained dog. Bar diagram on *top left* indicates that the relative density of dog cardiomyocytes with HCN4 immunolabelling obtained from the trained group (n=30 cells/6 dogs) is significantly increased compared to that measured in the sedentary group (n=30 cells/6 dogs). Original immunofluorescent images are shown on the *left*. *Bottom* panels indicate lack of effect of chronic training on the relative density of dog cardiomyocytes with HCN1 and HCN2 immunolabelling (n=30 – 30 cells/6 – 6 dogs for sedentary and trained groups, respectively). **Figure 7–Source Data 1** Effect of chronic training on HCN4 protein expression determined by immunocytochemical technique in sedentary and trained dogs. **Figure 7–Source Data 2** Effect of chronic training on HCN1 protein expression determined by immunocytochemical technique in sedentary and trained dogs. **Figure 7–Source Data 3** Effect of chronic training on HCN2 protein expression determined by immunocytochemical technique in sedentary and trained dogs.

## DISCUSSION

To the best of our knowledge this is the first study to provide comprehensive experimental data from a large animal model of sustained exercise regarding proarrhythmic electrophysiological remodeling and related increased proarrhythmic risk at the ventricular level. On the basis of the findings, we propose a novel hypothesis regarding the mechanism of SCD after endurance training. Specifically, we hypothesize that serious ventricular arrhythmias are related to the repolarization changes that develop after endurance training. This idea develops the work of Volders *at al.* [24] who presented experimental evidence that long lasting A-V ablation resulted in profound bradycardia, compensated cardiac hypertrophy, QTC, APD lengthening and enhanced spatial and temporal dispersion of repolarization. These changes were attributed to downregulation of several potassium currents [25] and associated with greatly increased ventricular arrhythmia susceptibility. We propose, therefore, that the mechanism of VF in our experimental animals, and that of elite athletes, have similar electrophysiological backgrounds. This is compatible with the experimental model of Volders *at al.* [24, 25] since the so called “athletes heart” shares some similarity with that of the compensated hypertrophy studied by Volders *at al.* [24]

The main findings of this study can therefore be summarized as following: 1, endurance training decreased heart rate both in whole animal and *in vitro* experiments. 2, Sustained training enhanced HCN4 protein expression, ventricular ectopic activity and fibrosis in canine ventricle. 3, After endurance training, moderate but significant APD lengthening and enhanced instability of cardiac repolarization were observed. These changes were associated with a decrease of the transient outward potassium current density. 4, Sustained exercise elevated the proarrhythmic risk subjected with electrical stimulation.

In elite athletes, there is an increased incidence of unexpected sudden death [6] the exact cause of which is poorly defined and as such is actually an open issue [7]. Our working hypothesis was that these relate to changes in ventricular repolarization and also in fibrosis which may occur in competitive athletes [26, 27]. However, to study the relevant mechanism is difficult for obvious reasons. Lengthening of PQ, QT_c_ intervals [28], bradycardia [28] and increased spatial and temporal dispersion of repolarization were reported in athletes reflected as enhanced T_e_-T_d_ interval and short term QT variability [29] but any putative link with increased incidence of death is unclear. There is, therefore, a pressing need for appropriate animal models to clarify the issue. So far, the majority of animal experiments in this field have been conducted in small animal models. However, these differ in important aspects of cardiac electrophysiology compared with humans thus limiting their translational value.

Electrophysiological remodelling, fibrosis, bradycardia and the possibility for enhanced risk of AF have been reported recently in rodents [4,8,11,12,30,31]. As far as we know, however, relatively scanty data has been obtained in large animal models of endurance training and proarrhythmic risk. Two chronic exercise models have been reported in rabbit [13, 32] but no such study has been reported on dog or other large animals. On the contrary, the available data on dogs show that acute [33] or chronic [34, 35] exercise exerts beneficial effects preventing ventricular fibrillation caused by acute ischaemia by delayed preconditioning [33] or by restoring the distorted parasymphathetic/sympathetic balance seen after myocardial infarction [34, 36]. The few available data on exercised dog regarding possible electrophysiological remodeling after sustained training originates from ECG recordings performed in long-distance sled race trained and corresponding control dogs. Although these studies showed that training significantly lengthens QRS and QT intervals [37, 38], these findings were not connected to arrhythmogenesis or electrophysiological remodeling at the cellular level.

In the present study, we found significant bradycardia after chronic exercise. Bradycardia in elite athletes and in endurance exercise animal models is a general and well established finding [39, 40] but its exact mechanism is still a matter of debate [12,39,41,42,43,44]. The most common explanation relates the bradycardia to enhanced vagal tone both in athletes [39] and in the corresponding animal studies [40, 41]. This latter studies generated some debate [9,12,45] which remains to be satisfactorily resolved. In addition, however, very recent publication by Mersica *at al.* [10] showed that in racehorses and mice after swimming-induced exercise, atrioventricular conduction slowing persisted after vegetative blockade further arguing for the role of I_f_ ion current downregulation and concomitant HCN4 ion channel remodelling as had been published earlier for mice and rat sinus node preparations [12]. The present data – at least partially - supports the observations of Souza at al [12] and Mersica *at al.* [10].

Accordingly, we show that a significant part of the sinus bradycardia still existed in isolated right atrial preparations. However, the enhanced rmsSD of the heart rate argues for an important contribution of enhanced vagal tone, as well. It should be emphasized, though, that the pacemaker function of the sinus node is complex and cannot be satisfactorily explained by the contribution of I_f_ only. Since activation of I_f_ largely occurs at voltages more negative than the maximal diastolic potential of sinus nodal cells the special importance of I_f_ as the sinus nodal main pacemaker current has even been questioned [46, 47]. In addition, other mechanisms based on calcium handling [48,49,50] or contribution of other ion channels were proposed [51,52,53] to explain cardiac pacemaking in the sinus node. Regardless of the nature of the bradycardia, slower heart rate itself would result in longer APD and enhanced dispersion of repolarization contributing to an increased substrate for arrhythmias. In addition, bradycardia which results in longer diastolic intervals would enhance the chances for spontaneous diastolic depolarization reaching the threshold for firing. In this context it should be emphasized that increased ventricular ectopic activity with concomitant elevation of HCN4 protein expression was also observed in the ventricle after chronic exercise in this study, similar to that described earlier in the failing heart [54, 55]. These findings argue in favor of ventricular ectopic activity serving as the potential trigger for arrhythmias. Further studies in this direction focusing on changes in intracellular calcium handling are now needed to clarify the issue.

The lengthening of repolarization in our experiments is a consistent observation and manifested as lengthened QT_c_ and APD in the whole animal and cellular measurements respectively. This finding was associated with increased STV of APD in the cellular measurements suggestive of enhanced spatial and temporal dispersion of repolarization, respectively. Similar repolarization changes were reported in the failing [56, 57] or hypertrophied heart [58]. These changes in combination with the mild degree of fibrosis would result in enhanced arrhythmia substrate providing somewhat wider vulnerable window for extrasystoles to provoke ventricular arrhythmias.

As an underlying mechanism of the observed repolarization lengthening in midmyocardial myocytes, only the decreased density of I_to_ was detected in the chronically exercised dogs compared to sedentary animals. It cannot, however, be excluded other currents, not examined in this study, could have a role. The electrophysiological response to exercise in different models is variable, not least because of the large variety of ion channel expression patterns that differ between species and tissue types. This further suggests the importance of studies in large animal models that better reflect the human condition. In a previous study from our group, it was found that inhibition of I_to_ in dog subepicardial muscle significantly lengthened APD [59]. Similar results i.e. APD lengthening associated with reduced I_to_ were the most consistent findings published in failing dog [56] or human [57] hearts. In mice atrial cells, however, no changes of I_to_ and I_Kur_ were noticed but I_Ca_ was upregulated after swim-induced chronic exercise [4]. In the recent report of Mesirca *at al.* [10] it was found that chronic swim-induced exercise in mice downregulated I_f_, I_Ca_ and I_Kr_ in atrioventricular nodal cells. Similarly in racehorses, Cav1.2 and HCN4 protein expression in atrioventricular nodal tissue was less than in sedentary controls [10]. In the present study, however, canine ventricular myocytes exhibited no change in either I_Ca_ and I_Kr_ after chronic exercise. The lack of repolarization lengthening in dog subendocardial papillary muscle preparation in this study would reflect the relatively weak [22, 23] I_to_ reported earlier in these preparations.

The smaller current density of Ito in exercised dogs in the present study seems not to be due to decreased expression of Kv4.3 alfa or KChIP2 beta accessory channel proteins since neither Western blots nor immunohistochemical measurements revealed differences in expression of these proteins in sedentary and trained hearts. Similar results i.e. decrease of I_to_ without changes of Kv4.2 and KChIP2 were reported earlier in chronically exercised rats [60]. Although Kv4.3 and KChIP2 are considered to be the most important proteins determining I_to_ in the dog [61], it has to be emphasized that the expression of other I_to_ accessory proteins such as Kvbeta1, Kvbeta2 [62, 63], I_Na_ beta1 [64], DPP6 [65, 66], DPP10 [67] may also affect I_to_ function [62], which were not investigated in the present study. In addition, PKA and PKC can modulate I_to_ [68, 69], and may also change after exercise.

Another consistent finding in this study was the enhanced fibrosis in dog ventricular muscle. This result is not surprising since several previous animal studies have reported similar findings after chronic endurance exercise [4, 8]. Since in our working hypothesis we focused on possible repolarization abnormalities to the molecular mechanism of increased fibrosis was not investigated in the present study. However earlier work by others investigated this issue in a great detail in mice and rat [4,11,30].

### Possible limitations

This work has some limitations. Athlete’s heart-associated arrhythmias including VF have very low incidence and the underlying cardiac electrophysiological alterations are relatively modest. In addition due to limited capacity of our lab in this study our investigation had to be focused on major transmembrane ionic currents but other cellular mechanisms like intracellular calcium handling or calcium dependent chloride, potassium and I_f_ pacemaker currents were not investigated The same was true for the molecular biological mechanism of the enhanced fibrosis. Therefore, to mimic these changes in experimental animal models require several months of chronic experimentations with relatively high number of animals. This is not particularly difficult with small animals like mice, or rat but more complicated with large animals like the dog. Such dog experiments are not only costly but more importantly require substantial facilities and trained personnel. Although the translational value of dog experiments is much [70] better than that of mice or rat, there are still known important electrophysiological differences regarding the repolarization reserve between the dog and human heart [16]. Taking into consideration that the maximal number of animals at each training session is usually less than possible with rodent studies the advantage of the dog experiments itself is also its limitation. Also, athlete’s heart-related arrhythmias and SCD cannot be attributed only to compensated cardiac hypertrophy but rather to combination of several coexisting factors such as HCM, drugs, doping, hypokalemia etc. which have not been investigated in the present study.

## Conclusion

In conclusion, we propose that after endurance training in elite athletes mild repolarization lengthening and enhanced repolarization instability occur which associate with mild ventricular fibrosis. These changes favor arrhythmia in the ventricle and in the presence of enhanced ectopic activity as a possible trigger they contribute to the very rare incidence of serious arrhythmias reported occasionally. However, it is essential to emphasize that the observed changes in this study are moderate as part of a normal adaptation to enhanced demand. In addition, the evidence regarding the beneficial influence of sport on health including the cardiovascular system is dominant. Therefore, we do not want to suggest at all that exercise even at a competitive level is harmful by eliciting ventricular arrhythmias in the normal heart. However, in certain individuals, or situations where repolarization reserve is impaired due to hidden diseases like HCM, LQT syndrome, diabetes or electrolyte disturbances, doping or otherwise harmless medications, endurance training can be an additional possible risk factor which should be taken into account to prevent possible complications in competitive sport.

## ACKNOWLEDGEMENTS

This work was supported by the National Research Development and Innovation Office (NKFIH K 135464 to AV, NKFIH PD-125402 and FK-129117 to NN, NKFIH K 128851 to IB, SNN-134497 to VV and GINOP-2.3.2.-15-2016-00047 and TKP2021-EGA-32), the Ministry of Human Capacities Hungary (20391 3/2018/FEKUSTRAT and EFOP-3.6.2-16-2017-00006), the UNKP-20-5-SZTE-165, János Bolyai Research Scholarship of the Hungarian Academy of Sciences (to NN), the Eötvös Loránd Research Network and by the Albert Szent-Györgyi Medical School institutional grant (SZTE ÁOK-KKA 2021 to LV).

## DATA AVAILABILITY

All data generated or analysed during this study are included in the manuscript and supporting file; Source Data files have been provided for Figures 1 - 7 and Table 1.

